# Quantification of biomolecular dynamics inside real and synthetic nuclear pore complexes using time-resolved atomic force microscopy

**DOI:** 10.1101/592303

**Authors:** George J. Stanley, Bernice Akpinar, Qi Shen, Patrick D. Ellis Fisher, C. Patrick Lusk, Chenxiang Lin, Bart W. Hoogenboom

## Abstract

Over the past decades, atomic force microscopy (AFM) has emerged as an increasingly powerful tool to study the dynamics of biomolecules at nanometre length scales. However, the more stochastic the nature of such biomolecular dynamics, the harder it becomes to distinguish them from AFM measurement noise. Rapid, stochastic dynamics are inherent to biological systems comprising intrinsically disordered proteins. One role of such proteins is in the formation of the transport barrier of the nuclear pore complex (NPC): the selective gateway for macromolecular traffic entering or exiting the nucleus. Here, we use AFM to observe the dynamics of intrinsically disordered proteins from two systems: the transport barrier of native NPCs, and the transport barrier of a mimetic NPC made using a DNA origami scaffold. Analysing data recorded with 50-200 ms temporal resolution, we highlight the importance of drift correction and appropriate baseline measurements in such experiments. In addition, we describe an auto-correlation analysis to quantify time scales of observed dynamics and to assess their veracity — an analysis protocol that lends itself to the quantification of stochastic fluctuations in other biomolecular systems. The results reveal the surprisingly slow rate of stochastic, collective transitions inside mimetic NPCs, highlighting the importance of FG-nup cohesive interactions.

## Introduction

With its ability to resolve biomolecules in aqueous solution at a spatial resolution of ~1 nm [1], AFM has a unique potential to observe dynamic biological processes at the single-molecule level. Thanks to technological advances in microscope hardware, cantilevers, and imaging modes, this high spatial resolution has become compatible with a temporal resolution on the order of 100 milliseconds per frame (often denoted as high-speed AFM, or HS-AFM) [2], or 0.5-1 millisecond for monitoring individual scan lines [3–5]. Over the years, AFM has allowed real-time visualization of diverse biological systems, from the binding/unbinding of GroEL molecules [3], to molecular motors walking along actin filaments [6], DNA cleaving by Cas9 [7], and a range of protein assembly processes [4,8–10] (amongst many other applications [2]).

When the observed dynamics are progressive in nature and have a magnitude well above the noise floor of the AFM, they can be unambiguously associated with molecular motion, binding, or conformational change. However, at molecular length scales, such dynamics invariably contain a stochastic component, and when the magnitude of these dynamics is ≲ 1 nm, they are not trivial to distinguish from AFM measurement noise or feedback errors that may arise when tracing molecular contours at high scan speeds. Such stochastic dynamics are arguably most pronounced for natively unfolded and intrinsically disordered proteins [11]. Unlike structured proteins, intrinsically disordered proteins have low sequence complexity and fluctuate rapidly over an ensemble of conformations ranging from extended statistical coils to more collapsed globules. Such proteins play a central role in a number of cellular processes: one important example being the selective filtering of macromolecules between the nucleus and the cytoplasm [12–14].

A large, proteinaceous self-assembly, known as the nuclear pore complex (NPC), controls this flow of materials into and out of the nucleus. The NPC has a cylindrical scaffold that perforates the nuclear envelope, creating a central channel of ~40 nm in diameter (see **Figure 1A and B**, schematics). This channel anchors intrinsically disordered proteins enriched with multiple phenylalanine-glycine repeats, known as FG-nucleoporins (or FG-nups), which, in combination with soluble proteins (nuclear transport receptors; or NTRs), collectively form a selective barrier to nucleocytoplasmic transport [15,16]. Exactly how they do this remains a contentious topic of debate [14].

**Figure 1.**
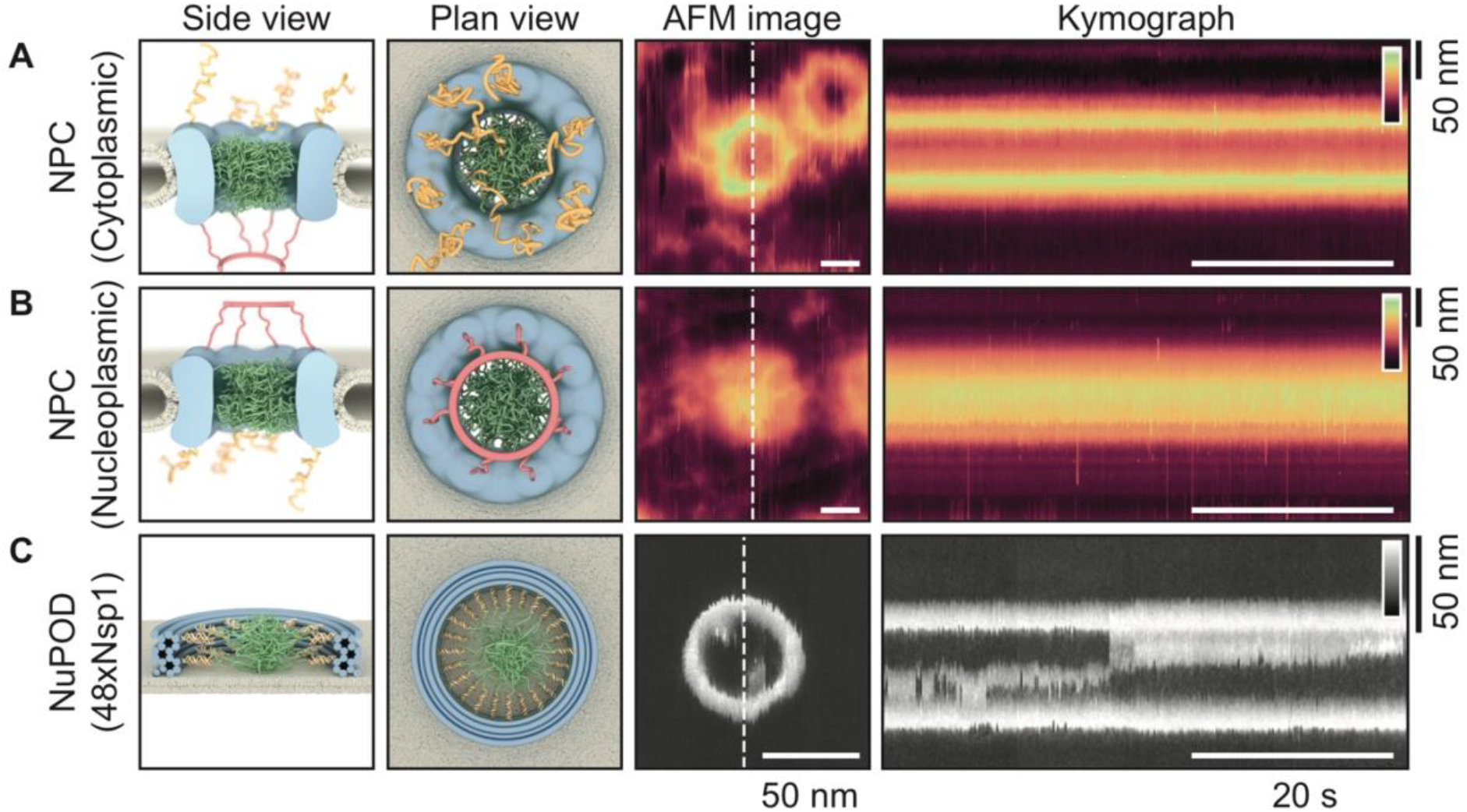
Producing kymographs from *Xenopus laevis* oocyte NPCs and from NuPODs. Schematics (side and top views) and representative AFM images of NPCs with the cytoplasmic **(A)** and nucleoplasmic **(B)** side facing the AFM tip, and of a DNA-origami based NPC mimic (‘NuPOD’ [5]; **C**) that contains 48 Nsp1 proteins, resting on a supported lipid bilayer. The NPC is asymmetrical when viewed through the mirror plane of the nuclear envelope: from the cytoplasmic face, cytoplasmic filaments are seen (yellow filaments); and from the nucleoplasmic face, the nuclear basket protrudes (red filaments). The disordered FG-nups are shown in green **(A-C)**. Kymographs are obtained by recording the same scan line across the pore (white-dashed lines in AFM images) as a function of time (horizontal axis of the kymographs). Note the differences in scale bars between the NPC and NuPOD data. Height scales (insets in kymographs): 80 nm for the NPCs, 20 nm for the NuPOD.

AFM has proven itself a useful tool for probing the dynamics of FG-nups inside real NPCs, with high-speed experiments potentially revealing dynamic movement of individual FG-nups at the cytoplasmic periphery of the pore [15,16], and high-resolution studies revealing many different metastable collective morphologies of FG-nups and soluble NTRs inside the central channel [17]. Both of these results support the notion that the “cohesivity” of the FG-nups is tuned such that the energetics of stable FG-nup morphologies lie near transition states, as has been postulated in computational studies [18–20]. One of the caveats with these experimental results, however, is that the exact chemical composition of the NPC’s transport barrier is ill-defined. Structural studies indicate that, by mass, the FG-nups only comprise (approximately) one third of the material inside the channel — the majority of the pore lumen is filled with NTRs and cargo molecules (*i.e.* macromolecules bound with NTRs) caught in transit [21]. Therefore, to better understand the behaviour and function of the FG-nups within this complex system, mimetic NPCs made using DNA origami have recently been created [5,22].

The NuPOD (<u>Nu</u>cleo<u>P</u>orins <u>O</u>rganised on <u>D</u>NA) system was designed to mimic the dimensions of the NPC’s central transport channel, and provide a platform for studying the behaviour of pure FG-nups in the cylindrical geometry [5]. This system has been used to accommodate two different S. *cerevisiae* FG-nups; Nup100, which is a cohesive GLFG-repeat nucleoporin; and Nsp1, a less cohesive FxFG-repeat nup. NuPODs therefore house purely the disordered FG-nups, of a known composition and grafting density (concentration).

In this study, we use AFM to probe the dynamic behaviour of disordered FG-nups across the transport barrier of both real NPCs, and mimetic NPCs (NuPODs) containing two different types of purified FG-nup domain (Nsp1 and Nup100). We present analysis, procedures, and criteria to distinguish between the stochastic dynamics of these proteins (often on length scales of ≲ 1 nm), and that of the (also largely stochastic) AFM measurement noise. By applying a drift-correction routine, and a subsequent auto-correlation analysis to the pixel heights with time, we are able to quantify both the time-scale of any dynamic behaviour observed, and probe the effect of the tip-sample interaction force on the behaviour of the system — thus providing a more rigorous approach to interpreting HS-AFM data. In doing so, we reveal the time-scale of collective dynamics of the pure FG-nups inside the NuPOD system, thus further elucidating their precise role in nucleocytoplasmic transport. For simplicity and clarity of representation, and for enhanced temporal resolution, we here confine ourselves to kymographs acquired by recording single lines across the centre of the pores as a function of time (**Figure 1**) [3], but note that our analysis can also be extended to frame-by-frame recordings (**Supplementary Figure 1**).

## Results and Discussion

When acquiring AFM data, temporal resolution can be greatly increased by reducing the dimension of scanning to the fast-scan axis only (as was done in this study), thereby producing kymographs. It can be increased further by removing this dimension also and recording the height of a single pixel with time (this has been termed HS-AFM height spectroscopy [4]). However, to accurately measure local height fluctuations as a function of time, lateral drift in the experimental setup must be minimised. Even when using an AFM system with a closed-loop scanner (here: Dimension FastScan, Bruker), in which the sample position is continuously verified to ~1 nm accuracy, lateral drift still occurs due to scanner hysteresis and thermal equilibration. Therefore, it is only by recording data in at least one dimension spatially (thus rendering points of reference), and employing a drift-correction routine, that we can ensure we are recording the same point in space with time. A drift-correction protocol is most accurate when referenced to static objects with steep slopes or protrusions in the data, as this is where a given lateral displacement results in the largest change in the measured AFM signal (*i.e.*, the sample height). Here, all scan lines in a kymograph were aligned laterally by referencing them to the outer edges of the static (and presumably rigid) pore scaffold (see **Supplementary Figure 2**).

With the kymographs corrected for experimental drift, measured height variations for any given position in the kymograph are — ideally — attributed solely to fluctuations as a function of time. Consequently, the magnitude and time scale of these height fluctuations can be quantified for each (lateral) position by the auto-correlation factor (*R*) as a function of time-lag (*τ*), defined as:

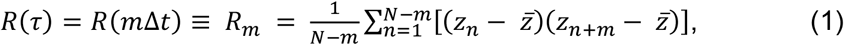

where *m* and *n* refer to line numbers in the kymograph, *N* is the total number of scan lines in the kymograph, and *Δt* is the time between consecutive scan lines. *Z_n_* denotes the pixel height (nm) of a given pixel as a function of time (*n*Δ*t*), and 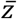 denotes the average pixel height for the (lateral) position under consideration. In brief, *R*(*τ*) quantifies the similarity in height values for a given pixel (in nm^2^) at a certain time-lag. It can detect a periodic signal shrouded by noise, and the persistence time of stochastically fluctuating signals (see **Supplementary Figure 3** for the *R*(*τ*) outputs from various simulated pixel fluctuations). Furthermore, it can be determined for each pixel in a kymograph, and the resultant *R*(*τ*) outputs can be represented as a heatmap as a function of both position (nm) and time-lag (*τ*), with red showing greater correlation and blue representing no correlation (see **Figure 2**).

**Figure 2.**
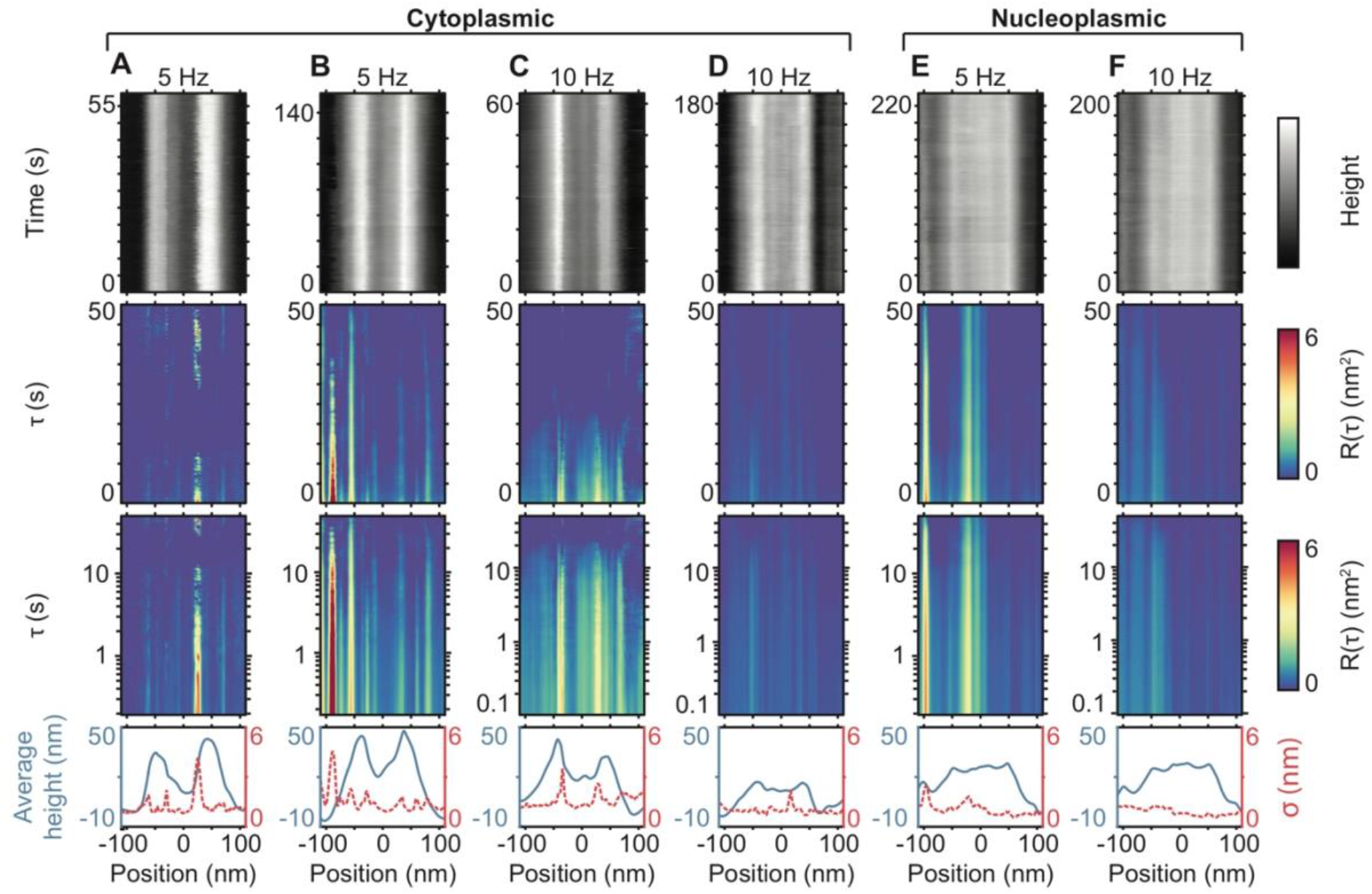
Kymographs and auto-correlation analysis of NPCs. **(A-D)** Kymographs of NPCs at the cytoplasmic face (top row), recorded as explained in **Figure 1**, but with time on the vertical axis (recorded at line frequencies of 5 and 10 Hz). The concomitant autocorrelation factor, *R*, is displayed as a function of position with respect to the pore centre and as a function of time-lag, *τ*, with *τ*plotted on linear (second row), and logarithmic (third row) scales. The average height profiles (blue) and respective standard deviations (red), are plotted in the final row. **(E and F)** Same as for **(A-D)** but for the nucleoplasmic face of NPCs. Kymograph height scales: 50 nm **(A and C)**, 60 nm **(B)**, 25 nm **(D)**, 40 nm **(E and F)**. **(A-D)** show data each from different NPCs; **(E and F)** is the same NPC scanned at 5 and 10 Hz.

When this analysis is applied to kymographs produced from the cytoplasmic and nucleoplasmic sides of intact NPCs (**Figure 2**), at different scan speeds, no significant fluctuations are detected in the positions of the central transport channel (i.e., around the 0 nm position). That is, the recorded fluctuations for positions occupied by the disordered FG-nups are not distinguishable to those at the positions of the nuclear envelope (a double membrane and lamin filaments, here supported by a glass substrate [17]). It remains possible that FG-nup dynamics inside the central channel yield Ångström-scale height variations (as have previously been reported for the cytoplasmic face of the transport channel of NPCs [15,16]), but in our experiments, based on a similar sample preparation, these fluctuations do not exceed the experimental noise floor. This is also apparent from the standard deviation for each position in the kymograph (**Figure 2**, bottom row).

However, significant fluctuations are detected — in both the *R*(*τ*) heatmaps and standard deviation plots — at positions that coincide with the edges of the NPC scaffold structure. At these edges, lateral drift most strongly affects the detected height (in spite of the applied drift correction, the observed fluctuations at the outer NPC scaffold tend to be larger than the fluctuations detected at the pore centre); and the height is more prone to tracking/feedback errors as the AFM tip traces protruding features. Hence, we attribute these fluctuations to artefacts of the AFM imaging process.

Similar artefactual fluctuations are observed when the analysis is applied to the empty pore scaffolds based on the biomimetic NuPOD system (**Figure 3A** and **Supplementary Figure 4**). By contrast, when the NuPODs contain grafted FG-nups, fluctuations are detected in the pore centre that greatly exceed those observed at the same positions in the empty NuPODs, or at the background supported lipid bilayer. This is here illustrated for NuPODs each containing a maximum of 48 copies of one of two types of FG-nup domains derived from the S. *cerevisiae* (yeast) NPC (**Figure 3B-G**; see Ref. [5] for further details). If 48 Nsp1 molecules are present (**Figure 3B-D**), fluctuating clumps inside the central channel are observed in the kymographs. At both 5 and 10 Hz (**Figure 3B and C**, respectively), the autocorrelation heatmaps show correlation up to ~10 s; at 20 Hz, little significant correlation is detected beyond 1 s (see **Figure 3D**, logarithmic plot). A similar pattern is observed for NuPODs containing 48 Nup100 molecules (**Figure 3E-G**): at 5 and 10 Hz, correlation is observed up to ~10 s, but at 20 Hz, this appears to be reduced. Furthermore, for all *R*(*τ*) heatmaps, no significant periodicity is detected — all heatmaps show signals more akin to stochastic fluctuating behaviour (see **Supplementary Figure 3C and D**).

**Figure 3.**
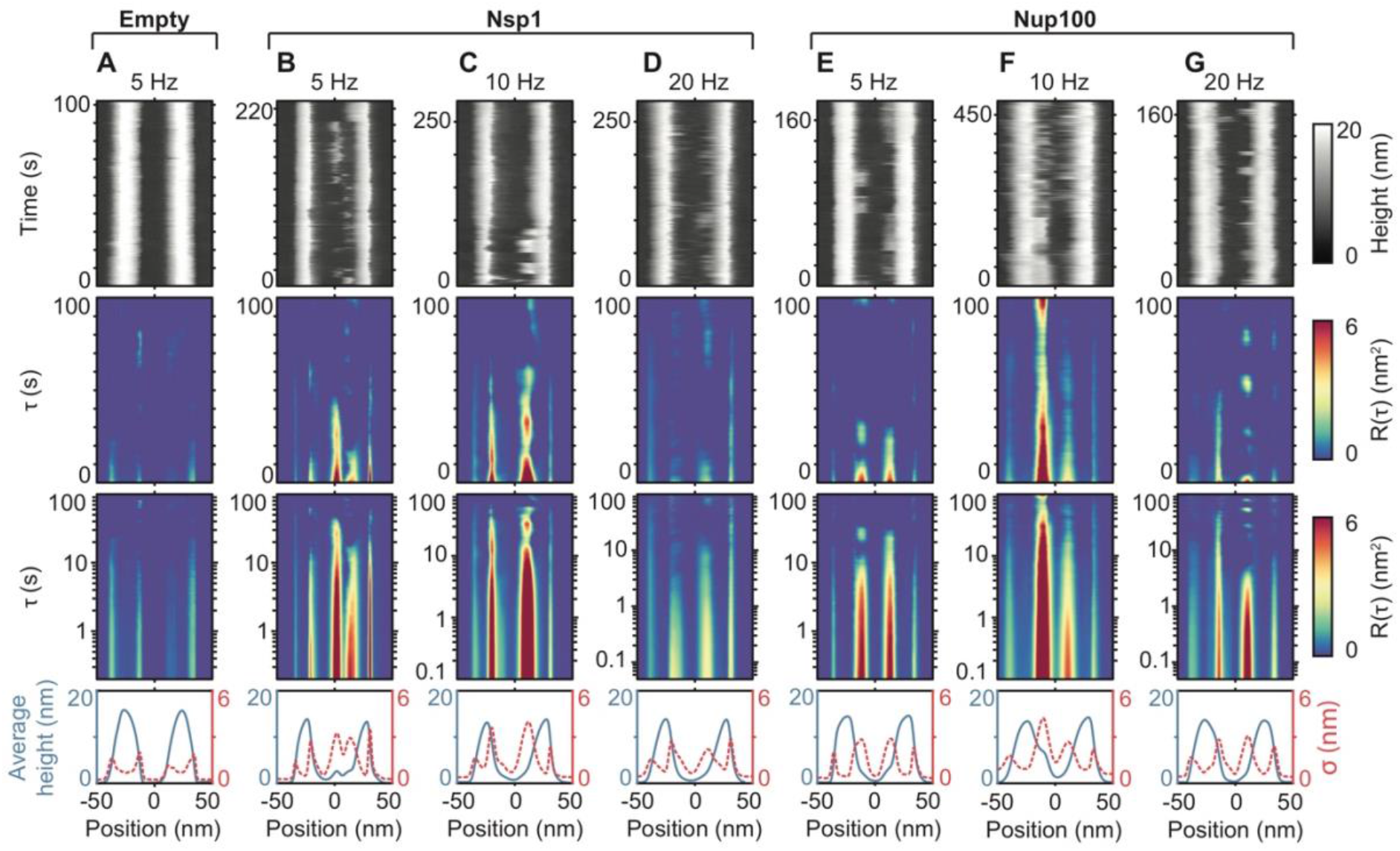
Kymographs and auto-correlation analysis of NuPODs. Repeat of kymograph data, auto-correlation analysis, and averaged height and standard deviation profiles, plotted as in ***Figure 2***, but on NuPOD NPC mimics. Data are shown for an empty NuPOD, *i.e.*, bare DNA scaffold without FG-nup domains **(A)**; for NuPODs containing 48 copies of Nsp1 **(B-D)**; and for NuPODs containing 48 copies of Nup100 **(E-G)**. The data of protein-containing NuPODs were recorded at scan line frequencies of 5, 10 and 20 Hz (200, 100 and 50 ms per scan line). **(A-D, and G)** show data each from different NuPODs; (E and F) show the same NuPOD scanned at 5 and 10 Hz.

A plausible interpretation of this result is that at higher scan speeds, more invasive feedback settings are required to track the NuPODs (data here were all obtained using Tapping Mode AFM, with the applied force minimised as much as possible without losing track of the pores). As a consequence, the flexible and mobile FG-nups may increasingly drop below the detection threshold of the AFM, perhaps in addition to being perturbed by the AFM tip. In this context, we note that it is not trivial to quantify the forces applied and energy dissipated into the sample [10]; and that it is nigh impossible to translate such forces and energies into plausible molecular deformations without knowledge of the dissipative mechanisms in the sample. To exclude that the observed dynamics are due to the invasiveness of the AFM measurements, the best approach remains to validate the observations at different scan speeds.

The results presented in **Figure 3** are from individual NuPODs. It is also possible to average over many auto-correlation heatmaps (produced from kymographs scanned at a given line rate), to build up a picture of the collective behaviour of each FG-nup, from many different NuPODs, over longer, aggregate recorded times. Furthermore, by cropping only a 30 nm window, which is smaller than the inner-diameter of the NuPOD (~46 nm [5]), it is ensured that only contributions from the FG-nups are considered. This eliminates any artefacts produced by tracking errors from the AFM tip scanning over the DNA scaffold structure. Next, by assuming the NuPODs display rotational symmetry, we can bin the data as a function of radial position, and average. This produces auto-correlation heatmaps as a function of radius, with 0 nm being the centre of the channel and 15 nm approaching the inner-wall of the scaffold. The NuPODs containing 48 Nsp1 molecules (**Figure 4A-C**) exhibit similar behaviour to that shown from the individual kymographs (**Figure 3B-D**), i.e., stochastic fluctuations with a decreasing persistence time as the scan rate increases from 5, to 10, to 20 Hz. A very similar picture is seen for the NuPODs containing 48 Nup100 molecules (**Figure 4D-F**). At 5 Hz, significant correlation is seen to up to ~10 s (**Figure 4D**); and at 10 Hz, it is seen to persist into the 1-10 s range (**Figure 4E**). At 20 Hz, however, the correlation does not persist much beyond 1 second (**Figure 4F**, logarithmic plot). All six heatmaps (**Figure 4A-F**) show stochastic, fluctuating behaviour (see **Supplementary Figure 3C and D**), and more correlation at the edges of the transport channels (*i.e.*, nearer radial values of 15 nm) as opposed to the centre (*i.e.* at 0 nm). Empty NuPOD controls confirm that these fluctuations are related to FG-nup fluctuations (see **Supplementary Figure 4**). Our observations indicate that the FG-nups spend more time clumped towards the inner-walls of the NuPODs, rather than in the middle of the transport channel; and that their rate of collective transitioning is recorded to increase with faster line scanning. Furthermore, at equal scan speeds, the more cohesive GLFG-repeat Nup100 tends to persist for longer before transitioning when compared to the FxFG-repeat Nsp1.

**Figure 4.**
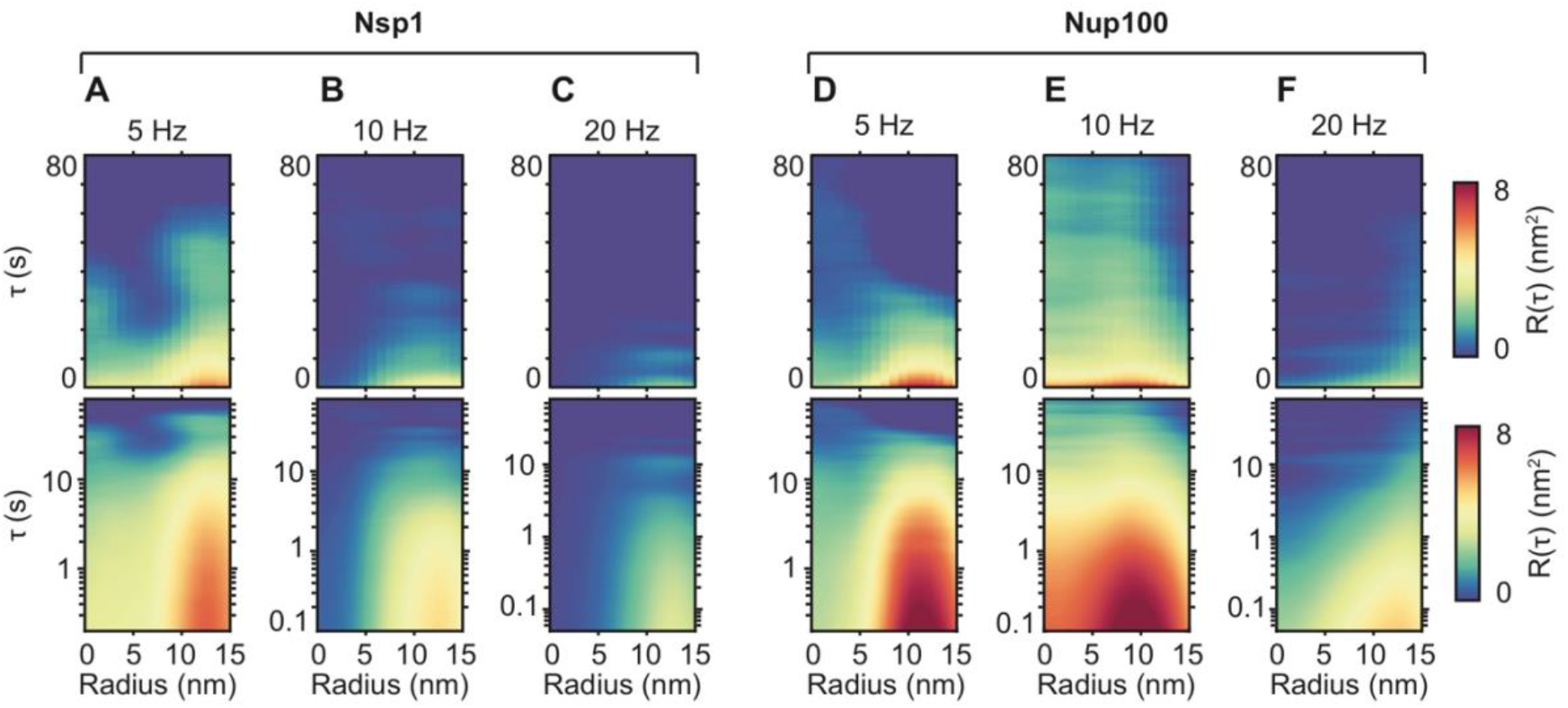
Averaged auto-correlation analysis of the pore channel in NuPODs. **(A-C)** *R*(*τ*) heatmaps averaged over *n* NuPODs, each containing 48 Nsp1 molecules — plotted as a function of radial position from the centre of the pore — scanned at line rates of 5, 10, and 20 Hz. *τ*is plotted on both linear (top row) and logarithmic scales (bottom row). **(D-F)** Same as for **(A-C)** but the NuPODs contain 48 Nup100 molecules. From **(A)** through to **(F)**: *n = 3, 4, 2, 3, 5, 3*; total duration of each kymograph = 739, 1050, 365, 586, 1907, 1664 s. It should be noted that one kymograph used in the averaging procedure to produce **(F)**, has previously been published as stand-alone data [5].

Taken together, these results show that, whilst no significant fluctuations are detected inside the native NPC (**Figure 2**), stochastic changes in height are detected for FG-nups inside the NuPOD system, persisting for up to ~10 s (when scanned at slower line rates: see **Figure 4A and D**). This can be further confirmed by plotting sample height at different locations in and around the pores as a function of time (**Figure 5**). For the NuPOD system, fluctuations inside the channel exceed those measured on and outside the pore scaffold by more than an order of magnitude (**Figure 5B and C**), whereas, for the native NPC, no noticeable difference in the magnitude of fluctuations is detected for different positions inside and outside the transport channel (**Figure 5A**). It is plausible that the discrepancy in behaviour observed between the native NPCs and NuPODs (i.e., non-fluctuating and fluctuating behaviour) is due to the presence/absence of endogenous NTRs (present in the real NPCs). In the preparation of native nuclear membranes from *Xenopus laevis* oocytes, the nuclei are repeatedly washed to remove cellular material — including cargoes and NTRs — that may be caught in transit. However, it is safe to assume that many NTRs and possibly cargo molecules bound to the FG-nups will not be removed by our washing steps [21]. In the case of the NuPODs, in the complete absence of NTRs, and in the symmetry of the pore geometry, there are many different possible metastable conformational states with small activation energy barriers between them — thus leading to the observed fluctuating behaviour. However, in the presence of multivalent NTRs (such as importin-β [23]), the FG-nups could wrap around these proteins, thereby becoming energetically trapped in a given conformation (as was observed for FG-nups at low grafting densities in planar films [24]). To this end, it would be interesting to acquire further kymograph data of NuPODs in the presence of NTRs — presumably, upon addition of importin-β, FG-nup fluctuations would stop. This experiment was attempted, but the addition of importin-β at physiological concentrations (~1 μM) disrupted the supported lipid bilayer in our experimental setup.

**Figure 5.**
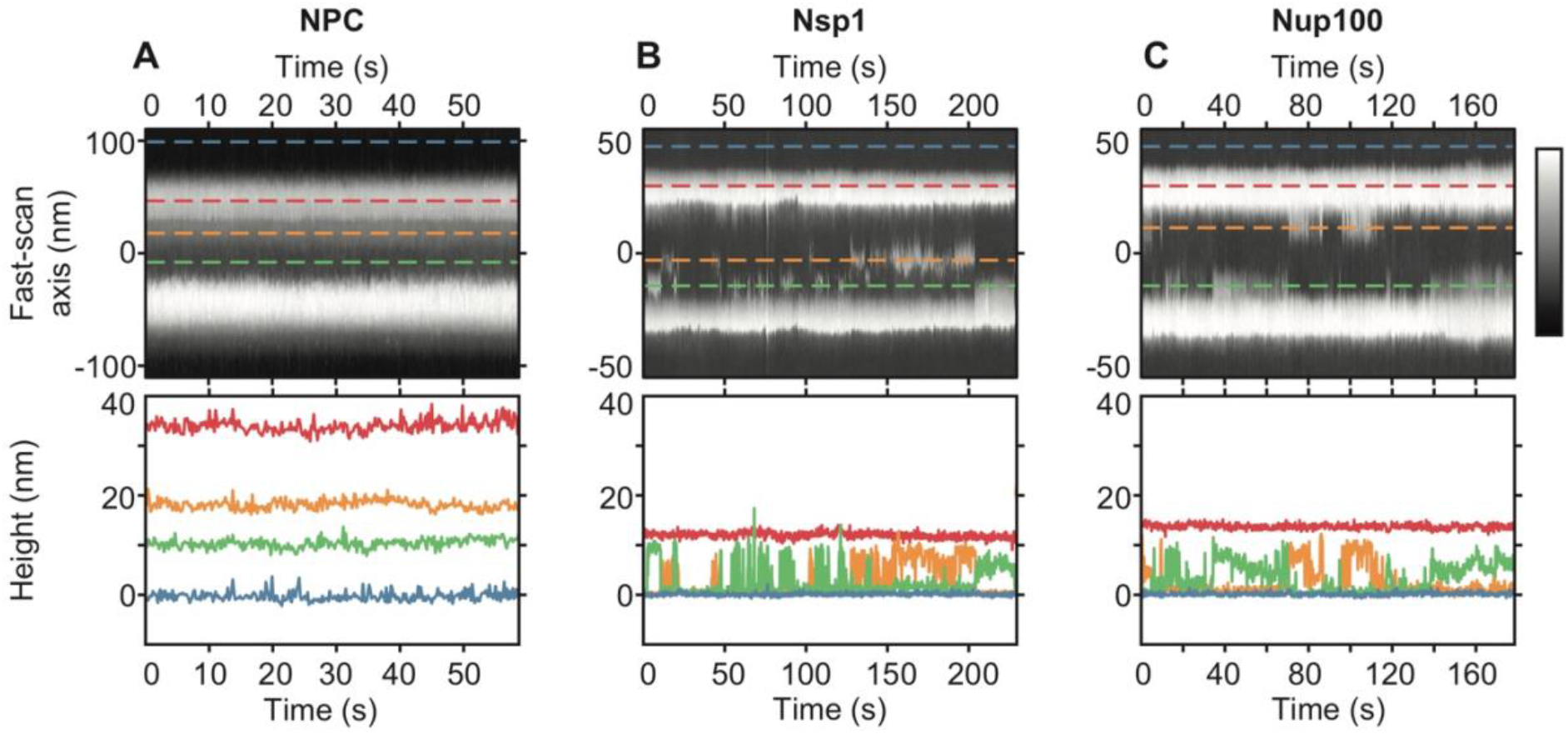
Kymographs and height fluctuations of NPCs and NuPODs. **(A)** Kymograph of a native NPC (cytoplasmic face) with the height-profiles of selected pixels shown underneath, scanned at 5 Hz. Blue-dashed line is the background nuclear envelope; red-dashed line is the NPC scaffold structure; orange- and green-dashed lines are different locations inside the transport channel. **(B and C)** Same as for **(A)** but of NuPODs containing 48 Nsp1 molecules **(B)** or 48 Nup100 molecules **(C)**. Blue-dashed lines are the background lipid bilayer; red-dashed lines are the DNA scaffold structure; orange- and green-dashed lines are different locations inside the channel. These kymographs are also printed in **Figure 2A**, and **Figure 3B and E**. Colour scales: 50 nm **(A)**, 20 nm **(B and C)**.

## Conclusion

Biologically, our results imply that pure FG-nups, grafted inside a cylinder, interact collectively to form transitioning clumps — thereby supporting the notion that their cohesive interactions are tuned such that stable morphologies lie near transition states [17–20]. Furthermore, we show a subtle divergence in behaviour between the FxFG-repeat Nsp1 and the more cohesive GLFG-repeat Nup100, with Nup100 persisting for longer times before transitioning (at comparable scan speeds; see **Figure 4**). Whilst this result is expected [25], it further emphasises the importance of cohesion — alongside molecular-scale dynamics — to the collective behaviour of the FG-nups. Nsp1, for example, which adopts a more extended state than Nup100 [25], is nevertheless shown to form clumps that can persist for several seconds before transitioning. In the native NPC, however, the static appearance of the FG-nups is probably due to the presence of other macromolecular contents (*i.e.*, NTRs and cargoes) that interact with the FG-nups and trap them in given morphologies [17,21].

More generally, our results highlight several criteria that can be applied to facilitate the interpretation of nanometre-scale fluctuations in time-resolved AFM data — especially of dynamic data obtained from biological systems exhibiting a stochastic, rather than progressive nature. Firstly, data must be background-corrected referenced to stable and flat areas of the sample (as common for AFM images) and — separately — corrected for lateral drift. Secondly, if fluctuations are observed in the vicinity of protruding features on the sample, additional controls are needed (e.g., the empty DNA scaffold control in **Figure 3** to rule out any artefactual nature). Thirdly, to rule out white noise, dynamic features should be observed to persist over multiple subsequent scan lines (in unfiltered data), such as observed here in the kymographs recorded on NuPODs, and further articulated by the autocorrelation analysis (**Figure 3**), demonstrating and quantifying the persistence time of the observed molecular dynamics. Fourthly, any such fluctuations should be significantly larger than fluctuations at other, presumably immobile positions on the sample (**Figure 5**). Finally, like any experimental method that probes individual molecules, AFM is an invasive technique: to identify the effect of the tip upon induced molecular dynamics, the observed movements should be shown to be relatively robust against variations in imaging parameters and/or speed (**Figure 4**).

## Materials and Methods

### NuPOD sample preparation and AFM imaging

#### NuPOD assembly

DNA origami structures and FG-domains conjugated to DNA were prepared as previously described [5]. These were stored at −20 and −80°C, respectively. The NucleoPorins Organised on DNA (NuPODs) were assembled by mixing DNA-labelled FG-nups with terminal MBP-tags in 7.5-fold excess (over the handle number) to DNA origami cylinders in NuPOD buffer (1x PBS, 0.1% Tween 20, 10% glycerol, 10mM MgCl2). After incubating at 37°C for 3 hrs the sample was cooled to room temperature and TEV-protease added to remove the terminal MBP-tag from the FG-nups now conjugated to the DNA rings. After a further incubation for 1.5 hours NuPODs were purified from excess protein by ratezonal centrifugation through a glycerol gradient.

#### Preparation of lipid vesicles

1, 2-dihexadecanoyl-sn-glycero-3-phosphocholine (DPPC) and dimethyldioctadecylammonium (bromide salt; DDAB) were purchased from Avanti Polar Lipids. DPPC and DDAB were mixed at a molar ratio of 3:1 in chloroform solvent. Lipid vesicles were prepared by the extrusion method as described elsewhere [5] and stored at 4°C for up to one week. Briefly, the chloroform was evaporated under a stream of nitrogen gas to yield a lipid film that was then re-dispersed with sonication in MilliQ water. The lipid vesicles were extruded through a 100 nm polycarbonate filter (GE Healthcare Lifesciences, Buckinghamshire, UK) and the extrusion process repeated at least 20 times to yield small unilamellar vesicles.

#### AFM sample preparation

2 μl of the lipid vesicles, along with 1 μl 1M MgCl_2_, 1 μl 1M CaCl_2_, and 16 μl of MilliQ water were deposited onto a freshly cleaved mica disc. The disc was placed in a humid chamber and heated to 65°C for 10 minutes to induce vesicle rupture. The sample was slowly cooled to room temperature over 20 minutes, to form a gel-phase supported lipid bilayer (SLB). Excess vesicles in the supernatant were removed by rinsing first with water and then exchanged to imaging buffer (10 mM PB 26 mM MgCl2, pH 7.0). The rinsing process was repeated three to five times to ensure a clean and uniform surface before addition of 2-4 μl of NuPODs (~2-5 nM) prior to imaging. The DNA origami scaffold is electrostatically adsorbed to the positively charged membrane.

#### AFM imaging

All AFM measurements were performed at room temperature in liquid. Images were obtained using a Dimension FastScan Bio AFM (Bruker) operated in Tapping mode. FastScan D (Bruker) cantilevers were used for all imaging with a resonance frequency of ~110 kHz, measured spring constant of ~0.15 Nm^-1^ and quality factor of ~2 in water. The force applied to the sample was minimized by setting the highest possible amplitude setpoint voltage, which was typically above 85% of free oscillation just close to the sample surface. Single line scanning experiments provided an enhanced time resolution whilst minimising disturbance to the NuPODs. For these experiments a single NuPOD was centred and the frame size decreased to 120 nm before disabling the slow-scan axis over the centre of the pore. Where possible, data was collected at 5, 10 and 20 Hz for each pore imaged.

### Nuclear envelope sample preparation and AFM imaging

#### Xenopus laevis *oocyte nuclear envelope preparation*

All experiments were conducted on *Xenopus laevis* oocyte NPCs. The oocytes were stored in modified Barth’s solution (88 mM NaCl, 15 mM Tris, 2.4 mM NaHCO_3_, 0.82 mM MgCl_2_, 1 mM KCl, 0.77 mM CaCl_2_, and U/100 μg penicillin/streptomycin, pH 7.4) at 4°C for a maximum of 3 d. The nuclei were isolated, and the nuclear envelopes prepared, in nuclear isolation medium (NIM) buffer (10 mM NaCl, 90 mM KCl, 10 mM MgCl_2_, 10 mM Tris, pH 7.4), for AFM imaging as previously described [17]. The nuclei are kept in buffer and on ice throughout the entire sample preparation; and no chemical fixation or detergent was used at any stage.

#### AFM imaging

All AFM measurements were performed at room temperature in import buffer (20 mM Hepes, 110 mM CH_3_COOK, 5 mM Mg(H_3_COO)_2_, 0.5 mM EGTA, pH 7.4). Kymographs were obtained using a Dimension FastScan Bio AFM (Bruker), using Tapping mode AFM. FastScan D (Bruker) cantilevers were used for all experiments, and the applied force was minimised (optimised) as described for the NuPODs. Images of the nuclear envelope were recorded to ascertain whether in contact with the cytoplasmic or nucleoplasmic face of the nuclear envelope [17]. A 300×300 nm image at 304 samples/line of a single NPC was recorded — this ensures capture of background nuclear envelope as well as the NPC. When the slow-scan axis is over the centre of the NPC (see **Figure 1A and B**, AFM images, white-dashed lines), it is disabled. A kymograph is now produced with height in the fast-scan axis and time in the slow-scan axis. The line rate is set to 5, 10, or 20 Hz, and the gains are optimised to best track the contours of the sample.

### Analysis protocols

All data analyses were done using MATLAB (MathWorks). The analysis code is accessible from GitHub: https://github.com/geostanley/AFM---Kymographs---Autocorrelation---DriftCorrection.

#### Concatenating kymographs

A sequence of kymographs recorded from one pore (NuPOD or NPC), at a given line rate, are loaded into MATLAB, and the kymograph width (nm), samples/line, and line rate (Hz) are entered manually. A 1^st^ order plane background subtraction is applied to each kymograph individually to flatten the data with respect to the background (either nuclear envelope or lipid bilayer). If the Down Scan Only feature during data capture was not enabled, the capture direction of the first kymograph must be entered (up or down). If the Down Scan Only feature was not enabled, the script vertically flips intermittent kymographs. If the Down Scan Only feature was enabled, this step is skipped. All images are then vertically concatenated. This gives one large kymograph with height in the fast-scan axis, and time (as captured chronologically) in the slow-scan axis. A 1^st^ order plane background subtraction is applied to the concatenated kymograph, again using the background nuclear envelope or lipid bilayer.

#### Drift-correction of kymographs

The concatenated kymograph is shown as a figure. The two outer edges of the scaffold (for either the NuPOD or NPC) are selected manually. Windows (usually ~10-20 nm in width) of kymograph data, centred around the two selected points, are cropped. The height data within these two windows are averaged to create a template. Each line (fast-scan axis) within the kymograph, at the same window positions, is then compared against the template using the sum of absolute differences (SAD) method: that is, the sum of absolute difference in height values between all relevant pixels is calculated, with a lower SAD score meaning greater correlation. Each line (or height array) is shifted in 1 nm intervals to both the left, and right, by half the window width. The SAD score is calculated at each position. Whichever position gives the lowest SAD score is considered to have the best correlation with the template, and the height array is moved to this position. If shifted, however, this results in missing data points at one end of the height array (if a row is shifted 6 nm to the left, for example, there will now be 6 nm of missing data points at the extreme right end of the array). As it is assumed that this region is either background lipid bilayer (for NuPODs) or background nuclear envelope (for NPCs), these pixels are filled in with random noise between the values of −1 and 1 nm. Schematics of this routine, for both the NPC and NuPOD, can be seen in **Supplementary Figure 2**.

#### Auto-correlation analysis

After drift-correction, an auto-correlation function is applied to the kymograph. This is defined as in Equation (1) in the Results and Discussion section. It should be noted that it is possible to normalise *R*(*τ*) values by dividing by the pixel’s variance (*σ*^2^). This would give *R*(*τ*) values between −1 and 1, at each time-lag, for each pixel, in which −1 is perfect anti-correlation and 1 is perfect correlation. However, to facilitate a quantitative comparison between experiments (in which all will have different *σ^2^* values for each pixel), this was not done. The *R*(*τ*) values are calculated for each pixel for *τ* = Δ*t* to *τ* = 100 s (or until the maximum time of the kymograph if recorded for less than 100 s), in increments of Δ*t*.

To average auto-correlation heatmaps, at a given line rate, from several NuPODs, the heatmaps must be aligned spatially. This was done by selecting the two inside edges of the scaffold structure as seen in the concomitant kymograph (akin to the drift-correction routine, in which both outer edges are selected). These two values are used to define the centre of the kymograph (and thereby the centre of their auto-correlation heatmaps). All autocorrelation heatmaps are then aligned by this point, and the *R*(*τ*) values 15 nm either side of this position are cropped. All heatmaps to be averaged are then scaled by their overall time contribution, and then summed. The data is then binned as a function of radial position, and averaged, to produce an auto-correlation heatmap as a function of radius, with 0 nm being the centre of the channel and 15 nm approaching the inner-wall of the scaffold (see **Figure 4**).

## Acknowledgements

The authors acknowledge Ariberto Fassati (University College London) for useful discussions; Alice Pyne (University College London) for training and advice on AFM; and the Yale Institute for Nanoscience and Quantum Engineering for use of AFM equipment. This work was funded by UK BBSRC and EPSRC studentships (BB/J014567/1 for G.J.S.; EP/L015277/1, EP/L504889/1 for B.A.); EPSRC equipment funding (EP/M028100/1); and by the NIH (R21GM109466 for Q.S. and C.P.L.).

## Author contributions

GJS designed and performed NPC experiments, developed analysis software, performed analysis, and wrote the manuscript. BA designed and performed NuPOD experiments, and wrote the manuscript. PDEF and QS prepared the DNA rings and purified protein constructs for NuPOD experiments. CPL and CL conceptualised the NuPOD system. BWH conceptualised the experiments and analysis, and wrote the manuscript. All authors reviewed and commented on the manuscript and its intellectual content.

## Supplementary Figures

**Supplementary Figure 1.**
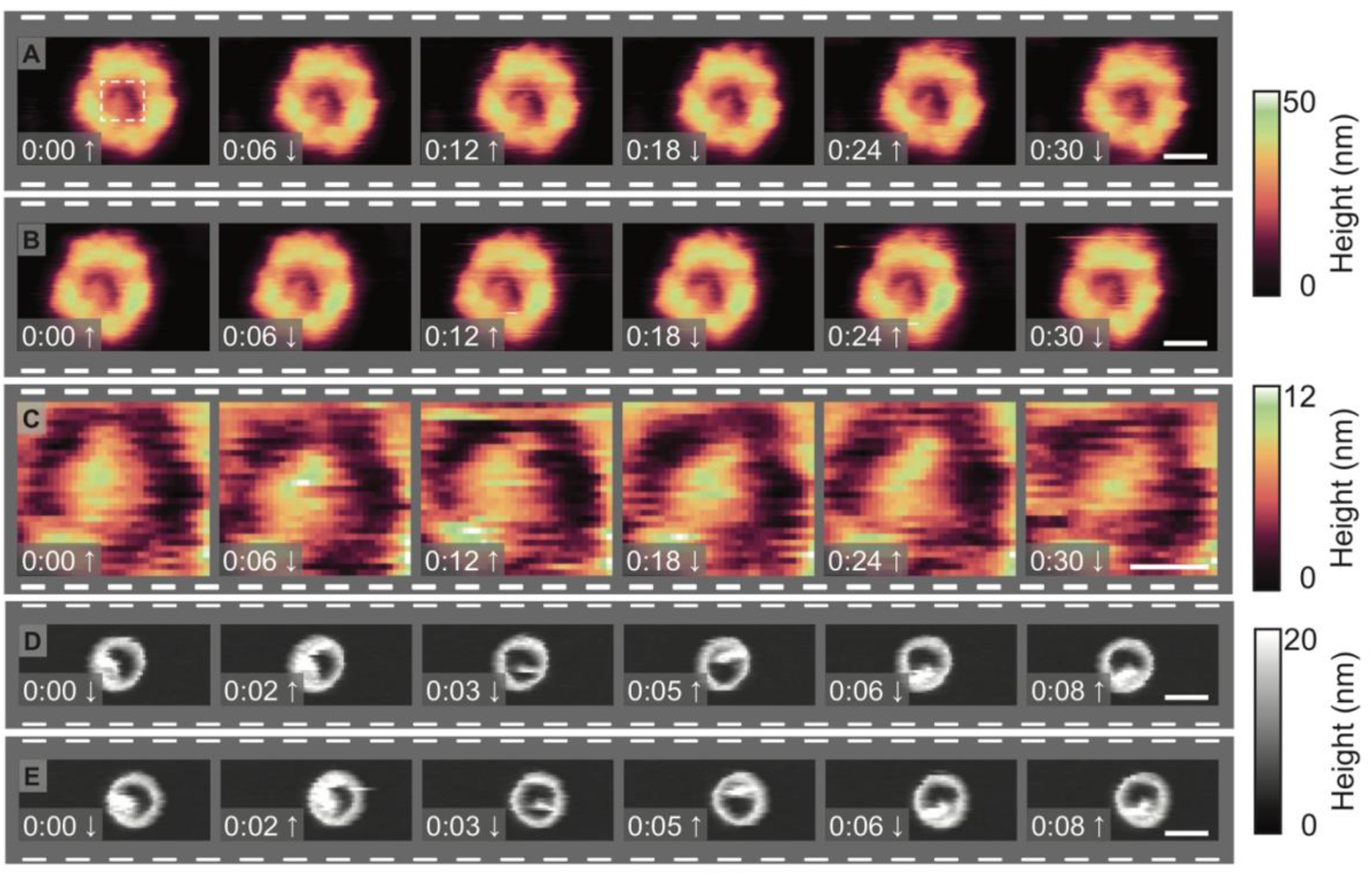
AFM video sequence of NPCs and NuPODs. **(A)** Consecutive AFM images of the cytoplasmic face of an NPC, recorded at ~6 s/frame. The NPC’s transport channel shows a stable protrusion. **(B)** Retrace images represent data acquired during the same scans as in **(A)**, but refer to the right-to-left line scans instead of the left-to-right line scans (‘trace’). These data faithfully reproduce the video sequence shown in **(A)**, but are slightly offset due to scanner hysteresis. **(C)** A digital zoom of the inside of the pore lumen, cropped from (**A**, see white-dashed box). A 2^nd^ order plane subtraction has been applied to each cropped image to better highlight any height differences within the pore lumen itself. **(D and E)** Trace and retrace images of a NuPOD containing 48 Nsp1 molecules, captured at ~1.6 frames/sec. Arrows indicate capture direction. These images represent snapshots extracted from data that have previously been published elsewhere in video format [5]. Scale bars: 50 nm **(A, B, D, E)**, 20 nm **(C)**.

**Supplementary Figure 2.**
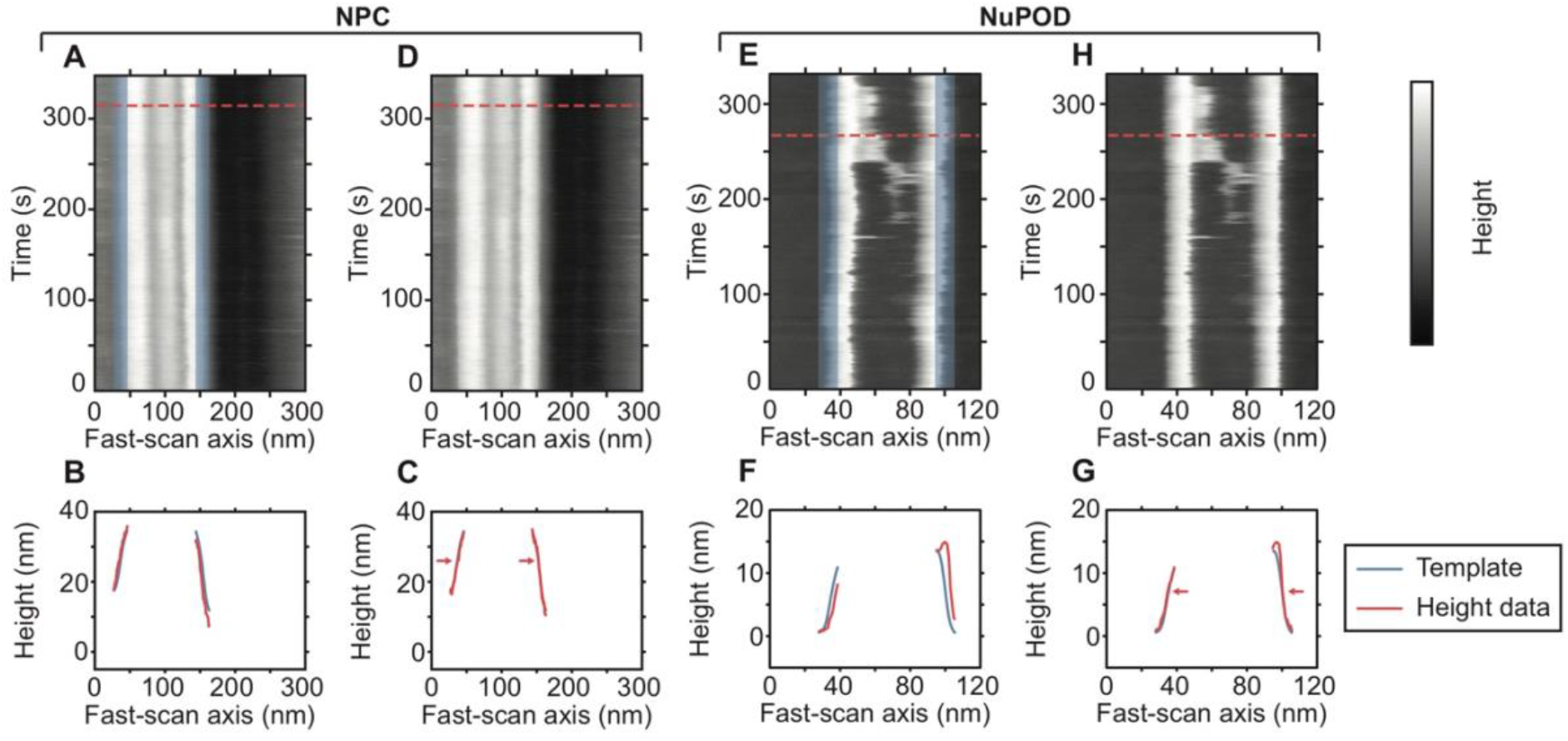
Drift correction of kymographs. **(A)** A kymograph of the cytoplasmic side of an NPC (scanned at 5 Hz). The outer edges of the pore scaffold are selected, and windows (typically of ~20 nm width) centred around these areas are cropped (blue, transparent regions). These regions are height averaged to produce a template (**B and C**, blue lines). Each line in the kymograph is then compared against this template within the window regions (red lines, **B**, corresponding to the red-dashed line in **A**). The data is then shifted in 1 nm intervals to both the left and right, for ~10 nm, and the sum of absolute difference of height values (SAD score) is calculated at each position. Whichever shift results in the best height correlation (or minimum SAD score), is used to correct that particular line **(C)**. **(D)** This operation is performed on every line in the kymograph to correct for lateral drift in our experiments. **(E-H)** Same as for **(A-D)** but demonstrated on a NuPOD containing 48 Nsp1 molecules.

**Supplementary Figure 3.**
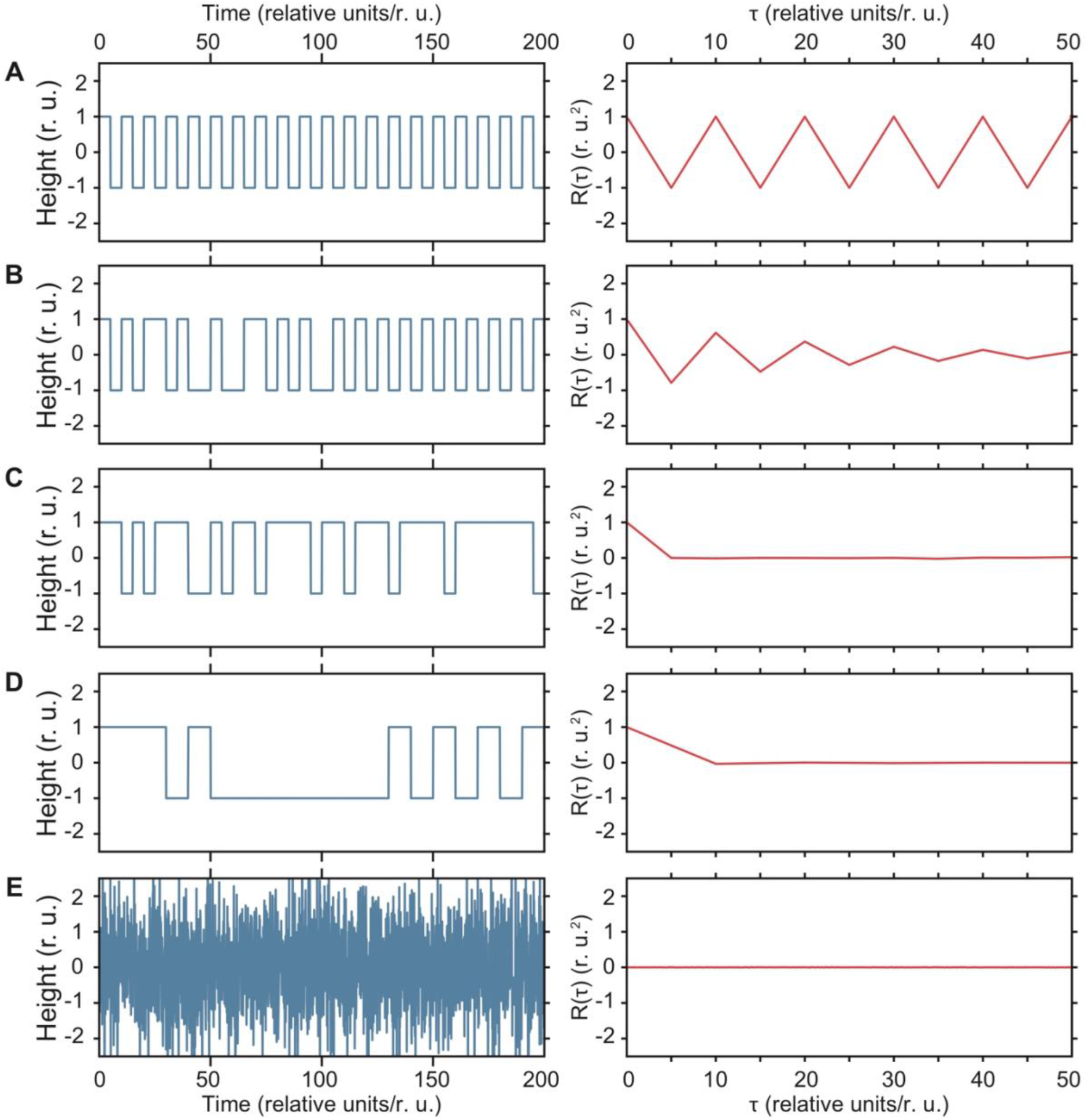
Interpreting the auto-correlation factor *R*(*τ*). **(A)** A simulated step function that changes between the values −1 and 1 at intervals of 5, therefore having a period of 10 (left panel), and calculated *R*(*τ*) (right panel). *R*(*τ*) is maximum for lag times, *τ*, that match integer multiples of the period of the original signal (*i.e.*, for *τ* = 0, 10, 20, *etc*.), indicating high correlation. By contrast, for half-integer multiples of the period (*τ* = 5, 15, 25, *etc*.), there is maximum anti-correlation. **(B and C)** Increased randomness in this function reduces long-scale auto-correlation, here shown when the flipping probability at each interval is reduced from 100% to 90% **(B)** and 50% **(C)**. In the case of **(C)**, no correlation is observed beyond a single half-period. **(D)** Same as **(C)**, but the interval length is increased from 5 to 10. Now correlation is observed up until τ = 10 instead of τ = 5. (E) For white noise, *R*(*τ*) is zero.

**Supplementary Figure 4.**
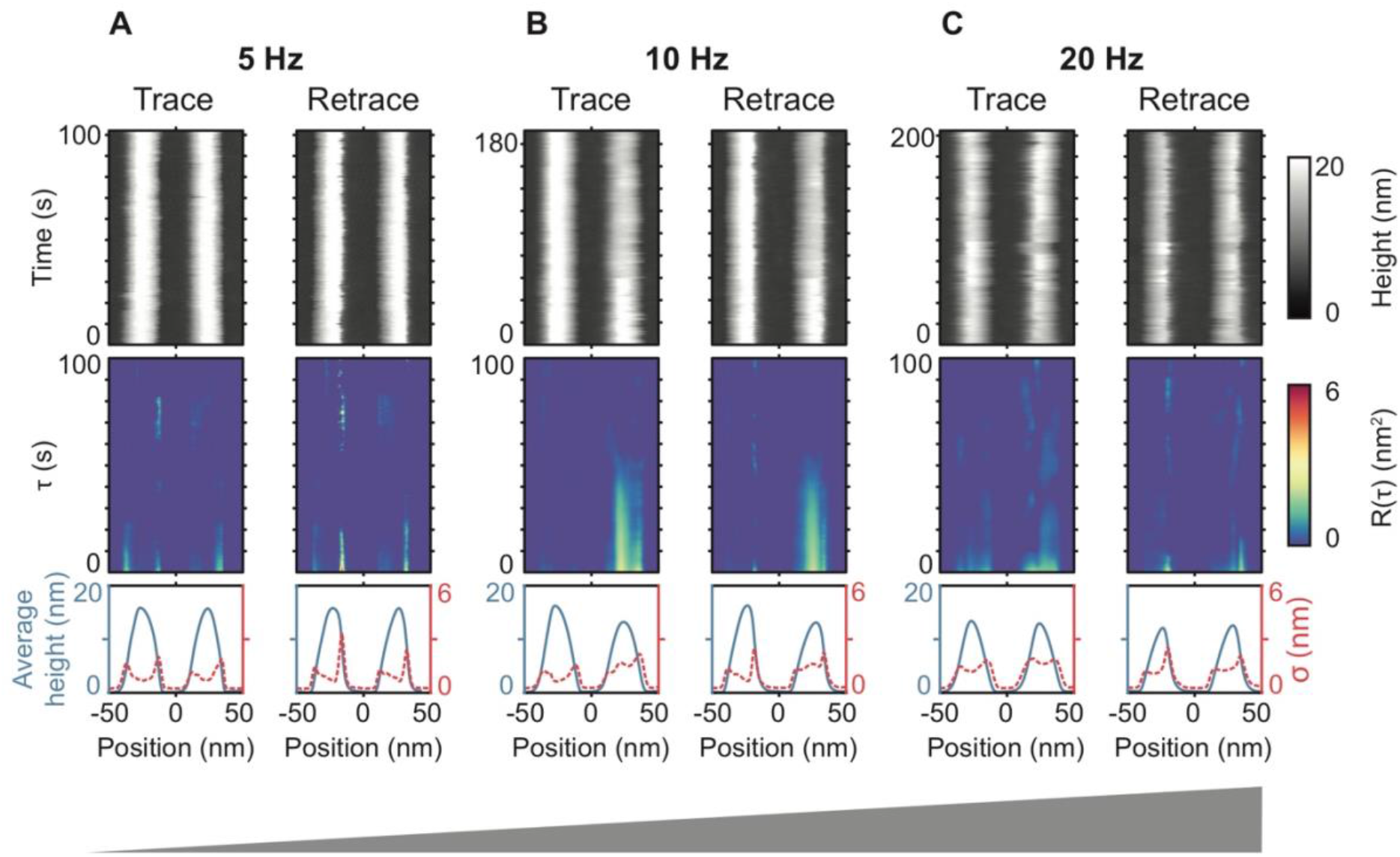
Kymographs and auto-correlation analysis of empty NuPODs. **(A-C)** Kymographs of NuPODs without FG-nups (top row), recorded at 5 **(A)**, 10 **(B)**, and 20 Hz **(C)**, showing both trace and retrace data. The light features are the DNA scaffold and the dark background the lipid bilayer. The auto-correlation heatmaps (second row) show background noise in the NuPOD lumens, and weak signatures of stochastic fluctuations at the edges of the DNA scaffold structure. Average height profiles of each pixel in the kymograph with time (bottom row, blue line), plotted with their respective standard deviation, *σ* (bottom row, red-dashed line), show maximum *σ*where the slope of the height profiles is steepest. Data shown in **(A)**, trace, is also printed in **Figure 3A**.

## References

1. Pyne A, Thompson R, Leung C, Roy D, Hoogenboom BW (2014) Single-Molecule Reconstruction of Oligonucleotide Secondary Structure by Atomic Force Microscopy. Small 10: 3257–3261.

2. Dufrêne YF, Ando T, Garcia R, Alsteens D, Martinez-Martin D, Engel A, Gerber C, Müller DJ (2017) Imaging modes of atomic force microscopy for application in molecular and cell biology. Nat Nanotechnol 12: 295–307.

3. Viani MB, Pietrasanta LI, Thompson JB, Chand A, Gebeshuber IC, Kindt JH, Richter M, Hansma HG, Hansma PK (2000) Probing protein-protein interactions in real time. Nat Struct Biol 7: 644–647.

4. Heath GR, Scheuring S (2018) High-speed AFM height spectroscopy reveals μs-dynamics of unlabeled biomolecules. Nat Commun 9: 4983.

5. Fisher PDE, Shen Q, Akpinar B, Davis LK, Chung KKH, Chung H, Baddeley D, Šarić A, Melia TJ, Hoogenboom BW, et al. (2018) A Programmable DNA Origami Platform for Organizing Intrinsically Disordered Nucleoporins within Nanopore Confinement. ACS Nano 12: 1508–1518.

6. Kodera N, Yamamoto D, Ishikawa R, Ando T (2010) Video imaging of walking myosin V by high-speed atomic force microscopy. Nature 468: 72–76.

7. Shibata M, Nishimasu H, Kodera N, Hirano S, Ando T, Uchihashi T, Nureki O (2017) Real-space and real-time dynamics of CRISPR-Cas9 visualized by high-speed atomic force microscopy. Nat Commun 8: 1430.

8. Chiaruttini N, Redondo-Morata L, Colom A, Humbert F, Lenz M, Scheuring S, Roux A (2015) Relaxation of Loaded ESCRT-III Spiral Springs Drives Membrane Deformation. Cell 163: 866–879.

9. Leung C, Hodel AW, Brennan AJ, Lukoyanova N, Tran S, House CM, Kondos SC, Whisstock JC, Dunstone MA, Trapani JA, et al. (2017) Real-time visualization of perforin nanopore assembly. Nat Nanotechnol 12: 467–473.

10. Nievergelt AP, Banterle N, Andany SH, Gönczy P, Fantner GE (2018) High-speed photothermal off-resonance atomic force microscopy reveals assembly routes of centriolar scaffold protein SAS-6. Nat Nanotechnol 13: 696–701.

11. Miyagi A, Tsunaka Y, Uchihashi T, Mayanagi K, Hirose S, Morikawa K, Ando T (2008) Visualization of Intrinsically Disordered Regions of Proteins by High-Speed Atomic Force Microscopy. ChemPhysChem 9: 1859–1866.

12. Stewart M (2007) Molecular mechanism of the nuclear protein import cycle. Nat Rev Mol Cell Biol 8: 195–208.

13. Wente SR, Rout MP (2010) The Nuclear Pore Complex and Nuclear Transport. Cold Spring Harb Perspect Biol 2: a000562.

14. Stanley GJ, Fassati A, Hoogenboom BW (2017) Biomechanics of the transport barrier in the nuclear pore complex. Semin Cell Dev Biol 68: 42–51.

15. Sakiyama Y, Mazur A, Kapinos LE, Lim RYH (2016) Spatiotemporal dynamics of the nuclear pore complex transport barrier resolved by high-speed atomic force microscopy. Nat Nanotechnol 11: 719–723.

16. Mohamed MS, Kobayashi A, Taoka A, Watanabe-Nakayama T, Kikuchi Y, Hazawa M, Minamoto T, Fukumori Y, Kodera N, Uchihashi T, et al. (2017) High-Speed Atomic Force Microscopy Reveals Loss of Nuclear Pore Resilience as a Dying Code in Colorectal Cancer Cells. ACS Nano 11: 5567–5578.

17. Stanley GJ, Fassati A, Hoogenboom BW (2018) Atomic force microscopy reveals structural variability amongst nuclear pore complexes. Life Sci Alliance 1: e201800142.

18. Osmanović D, Bailey J, Harker AH, Fassati A, Hoogenboom BW, Ford IJ (2012) Bistable collective behavior of polymers tethered in a nanopore. Phys Rev E 85: 061917.

19. Zahn R, Osmanović D, Ehret S, Araya Callis C, Frey S, Stewart M, You C, Görlich D, Hoogenboom BW, Richter RP (2016) A physical model describing the interaction of nuclear transport receptors with FG nucleoporin domain assemblies. Elife 5: e14119.

20. Vovk A, Gu C, Opferman MG, Kapinos LE, Lim RYH, Coalson RD, Jasnow D, Zilman A (2016) Simple biophysics underpins collective conformations of the intrinsically disordered proteins of the Nuclear Pore Complex. Elife 5: e10785.

21. Kim SJ, Fernandez-Martinez J, Nudelman I, Shi Y, Zhang W, Raveh B, Herricks T, Slaughter BD, Hogan JA, Upla P, et al. (2018) Integrative structure and functional anatomy of a nuclear pore complex. Nature 555: 475–482.

22. Ketterer P, Ananth AN, Laman Trip DS, Mishra A, Bertosin E, Ganji M, van der Torre J, Onck P, Dietz H, Dekker C (2018) DNA origami scaffold for studying intrinsically disordered proteins of the nuclear pore complex. Nat Commun 9: 902.

23. Isgro TA, Schulten K (2005) Binding Dynamics of Isolated Nucleoporin Repeat Regions to Importin-β. Structure 13: 1869–1879.

24. Lim RYH, Fahrenkrog B, Deng J, Aebi U (2007) Nanomechanical Basis of Selective Gating by the Nuclear Pore Complex. Science (80-) 318: 640–644.

25. Yamada J, Phillips JL, Patel S, Goldfien G, Calestagne-Morelli A, Huang H, Reza R, Acheson J, Krishnan V V, Newsam S, et al. (2010) A Bimodal Distribution of Two Distinct Categories of Intrinsically Disordered Structures with Separate Functions in FG Nucleoporins. Mol Cell Proteomics 9: 2205–2224.

